# O-glycosylation regulates plant developmental transitions downstream of miR156

**DOI:** 10.1101/618744

**Authors:** Krishna Vasant Mutanwad, Alexandra Baekelandt, Nicole Neumayer, Claudia Freitag, Isabella Zangl, Dirk Inzé, Doris Lucyshyn

**Affiliations:** Department of Applied Genetics and Cell Biology, University of Natural Resources and Life Sciences, Vienna, Muthgasse 18 1190 Vienna, Austria; Ghent University, Department of Plant Biotechnology and Bioinformatics, Technologiepark 71, 9052 Ghent, Belgium; VIB Center for Plant Systems Biology, Technologiepark 71, 9052 Ghent, Belgium

**Keywords:** *Arabidopsis thaliana*, O-glycosylation, plant development, phase change, flowering time, SPL

## Abstract

The timing of plant developmental transitions is decisive for reproductive success and thus tightly regulated. The transition from juvenile to adult vegetative and later to the reproductive phase is controlled by an endogenous pathway regulated by miR156, targeting the SQUAMOSA PROMOTER BINDING PROTEIN (SBP/SPL) family of transcription factors. SPLs regulate a number of developmental processes, such as trichome formation, leaf shape and floral transition. Such complex regulatory pathways often involve post-translational modifications (PTMs), integrating a range of internal and external signals. One of these PTMs is O-glycosylation, the attachment of a single monosaccharide to serine or threonine of nuclear and cytoplasmic proteins, which is found on a number of very diverse proteins. O-GlcNAcylation is the most common type of cytosolic O-glycosylation, but in plants also O-fucose modification occurs. Here we show that mutants defective in the O-fucosyltransferase SPINDLY (SPY) show accelerated developmental transitions. Genetic analysis shows that this effect is independent of miR156 levels, but partly dependent on functional SPLs. In a phenotyping analysis, we found that SPY and SPLs also control leaf growth, as loss of function mutants showed defects in cell expansion, while SPL9 also regulates cell division in rosette leaves. Moreover, SPLs interact directly with SPY and are O-glycosylated. Our results show that O-glycosylation is involved at several steps in the regulation of developmental transitions and organ growth in *Arabidopsis thaliana*.

## INTRODUCTION

During their life cycle, plants undergo developmental transitions by changing from the juvenile to adult vegetative and later the reproductive phase [1-3]. Each of these irreversible transitions is marked by specific morphological and developmental changes, and while they are governed by an internal developmental program, the timing is flexible and can be influenced by the environment. Both of these phase transitions are controlled by small RNAs, the families of miR156 and miR172. miR156 is encoded by eight genes, and there is at least partial redundancy with the closely related miR157, which is encoded by four genes [4]. miR156 directly targets a group of transcription factors within the family of SQUAMOSA PROMOTER BINDING PROTEINs (SBP/SPL), which then control both transitions by inducing a number of transcription factors regulating adult leaf traits, floral transition and floral meristem identity, as well as the expression of miR172 [5-14]. The family of SPLs contains 16 members, but only well as the expression of miR172 [5-14]. The family of SPLs contains 16 members, but only 10 of them are targeted by miR156. Interestingly, regulation occurs both by transcript cleavage as well as translation repression, reflecting the high level of complexity in the regulatory pathways of developmental transitions [4, 15]. Additionally, the decrease in miR156 expression throughout development is strongly influenced by metabolism: the accumulation of sugars during plant growth leads to a decrease in miR156 and thus an increase in miR172, which then additionally feeds back negatively on the expression of miR156 [16, 17]. Thereby as plants grow and build up biomass, the increasing availability of sugars leads to a release of the repression of SPL transcription factors and thus the developmental switch from the juvenile to adult and later reproductive phase. At the same time, organ growth is promoted by a transition from the proliferative phase characterized by cell divisions, to a phase where cell expansion is driving growth. This is regulated by GROWTH-REGULATING FACTORs (GRFs), a group of transcription factors regulated by miR396 [18-20]

The timing of transitions is extremely important for plants to ensure reproductive success. Thus, the transition from the vegetative to the reproductive phase is controlled not only by the internal developmental program described above, but also by seasonal cues and environmental conditions, such as day length, ambient temperature or light quality, which are integrated at several levels. Environmental factors, most importantly day length, are perceived in the leaves, where they induce the expression of the floral integrator *FLOWERING LOCUS T* (*FT*). FT is a small, mobile protein, moving through the vasculature to the shoot apical meristem, where it interacts with the transcription factor FLOWERING LOCUS D (FD) to induce flowering [21-23]. In the shoot apical meristem, the floral integrators SOC1 (SUPRESSOR OF OVEREXPRESSION OF CONSTANTS 1), LFY (LEAFY) and FT regulate the transcriptional network that underlies flowering time control, as reviewed in [24]. A role for gibberellins (GAs) in the induction of flowering time has already been suggested in the 1950s [25, 26], and research on the long-day plant *Arabidopsis thaliana* has shown that GA is necessary for floral induction under short photoperiods [27]; In long days, its effect is mostly masked by the much earlier responding photoperiod pathway [28]. Gibberellins have since been placed at several points within the flowering time regulatory network [11-13, 29-33], with *SOC1, LFY* and/or *SPLs* being direct targets. The effect of GA on flowering time is mediated by DELLA proteins, a small family of five transcriptional repressors that are negatively regulated by GA [34]. In the absence of GA, DELLA proteins bind to a number of transcription factors, thereby preventing their ability to bind their target genes [35-37]. The miR156-regulated aging pathway is independent of photoperiod, but is affected by gibberellins, as SPL15 is directly inhibited by the interaction with the DELLA protein RGA in the absence of GA [12].

O-glycosylation of nucleocytoplasmic proteins is an abundant post-translational modification, with diverse targets. In plants, two O-glycosyltransferases are described in this context, the O-GlcNAc Transferase (OGT) SECRET AGENT (SEC) and the Protein O-Fucosyltransferase (POFUT) SPINDLY (SPY) [38, 39]. These enzymes use UDP-GlcNAc (N-acetylglucosamine) and GDP-fucose, respectively, to transfer the single sugar moiety to serine or threonine residues on their target proteins. Among these targets are the GA signaling-repressing DELLA proteins as well as a number of other transcriptional regulators [38-40]. So far, only few examples of regulation by O-glycosylation have been characterized in plants. While *sec*-mutants show only very subtle phenotypes, *spy*-mutants show a range of developmental defects, and most of them have been explained by enhanced GA signaling [38, 39, 41-43]. One of the most prominent features of *spy*-mutants is early flowering. In long photoperiods, SPY acts together with GIGANTEA (GI) to repress expression of *CONSTANS* (*CO*) and *FT* [44]. SPY also strongly represses flowering in short photoperiods, which has been explained by its role in regulating GA signaling [41, 42].

Here, we present a further characterization of the role of SPY in the control of floral transitions and show an additional function of O-glycosylation in the regulation of the transition from juvenile to adult vegetative, and then to the reproductive phase. A combination of genetic and phenotypic analysis suggests a direct regulation of SPL-transcription factors downstream of miR156 by O-glycosylation. We also observed that leaf growth is strongly affected by SPY and SPLs at the level of cell expansion. While there was also an effect of SPL9 on cell division, we did not see an indication for control of cell division by glycosylation. We describe effects of protein O-glycosylation that are potentially independent of gibberellins, with redundancy between O-GlcNAc and O-fucose modification, but a much stronger effect of O-fucosylation.

## RESULTS

### The O-fucosyltransferase SPINDLY delays flowering in long and short photoperiods

In order to characterize the effect of O-glycosylation on flowering time regulation, we studied flowering time in O-glycosylation mutants. As strong *spy* alleles show severely reduced fertility [39, 41], we used the T-DNA-insertion line *spy-22* (SALK_090582) for all our experiments. This line showed strongly reduced expression of SPY (Fig.S1A-B) and was flowering early at 8.6 ±1.4 total rosette leaves (TRL), compared to 13.1 ± 1.1 in the wild-type Col-0 (Figure 1A-B, Table 1). The early flowering of *spy-22* was complemented by a *SPY::SPY:Flag* (*SPY:Flag*) construct (Fig.S1C-D). Recently, modest early flowering of *sec-5* was reported [45], which we also observed, with a very small difference to the wild-type (11.8 ± 1.6 TRL in *sec-5* compared to 13.1 ± 1.1 in Col-0; Figure 1A-B, Table 1). Accordingly, expression of the major floral integrator gene *FT* was up-regulated in *spy-22* but not *sec-5*, (Figure 1C). When grown in short photoperiods (8 h light / 16 h dark), where *FT* is not expressed, *spy-22* also displayed strong early flowering with 22.7 ± 0.5 TRL compared to 66.0 ± 7.9 in Col-0, which we did not observe in *sec-5*, flowering at 61 ± 5.0 TRL (Figure 1D-E, Table 1). Early flowering of *spy-22* was only partially suppressed by *ft-10*, resulting in a phenotype of *ft-10 spy-22* (17.4 ± 1.4 TRL) intermediate between *ft-10* (47.7 ± 2.9 TRL) and the wild-type Col-0 (14.6 ± 1.3 TRL), while *ft-10 sec-5* (46.2 ± 6.4 TRL) was comparable to *ft-10* (Figure 1F, Table 1). This confirms previously shown results with mutants in the L*er*-0 background [44], indicating that SPY regulates flowering by regulating *FT* and additionally in an *FT*-independent manner. This suggests that additional factors independent of the photoperiod pathway may be affected by SPY, while SEC appears to play a minor role. Therefore, we also generated crosses with the late flowering Col-0 FRI. This line carries an active FRIGIDA (FRI) allele introgressed from the ecotype Sf-2, and consequently expresses high levels of the floral repressor *FLOWERING LOCUS C* (*FLC*) leading to very low levels of *FT* expression and requirement of vernalization for floral induction [46, 47]. Late flowering of Col-0 FRI (74.2 ± 7.8 TRL) was strongly accelerated in *spy-22* FRI (26.0 ± 1.4 TRL, Figure 1G), but it was not affected by *sec-5* (data not shown). Levels of *FLC* expression were maintained high in that line, and *FT* was repressed (Figure 1H), suggesting that SPY represses flowering in Col-0 FRI independently and in addition of the vernalization pathway and FT. Thus, our data indicate that SPY represses floral transition only partly via FT and the photoperiod pathway. Additionally, SPY is necessary to maintain the vernalization requirement of Col-0 FRI.

**Table 1:** Rosette leaf numbers for graphs shown in Figure 1.

| | | Rosette leaves (average ± SD) | n |
| --- | --- | --- | --- |
| LD | Col-0 | 13.1 ± 1.1 | 51 |
| | spy-22<sup>a</sup> | 8.6 ± 1.4 | 46 |
| | sec-5<sup>a</sup> | 11.8 ± 1.6 | 49 |
| SD | Col-0 | 61.0 ± 6.1 | 33 |
| | spy-22 | 20.7 ± 3.1 | 33 |
| | sec-5 | 60.5 ± 6.7 | 30 |
| LD | | Rosette leaves (average ± SD) | n |
| | Col-0 | 14.6 ± 1.3 | 13 |
| | ft-10 | 47.6 ± 2.9 | 9 |
| | spy-22 | 7.1 ± 0.6 | 13 |
| | sec-5 | 14.9 ± 1.5 | 11 |
| | ft-10 spy-22 | 17.4 ± 1.3 | 11 |
| | ft-10 sec -5 | 46.2 ± 6.4 | 8 |
| | | Rosette leaves (average ± SD) | n |
| | Col-0 | 14.7 ± 1.3 | 13 |
| | spy-22 | 7.1 ± 0.7 | 13 |
| | FRI | 74.2 ± 7.8 | 12 |
| | FRI spy-22 | 26.0 ± 1.4 | 10 |

average ± SD is given
One-way ANOVA, Tukeys Multiple Comparison Test:
<sup>a</sup> significantly lower than Col-0 (\*\*\*) $p \leq 0.001$

**Figure 1.**
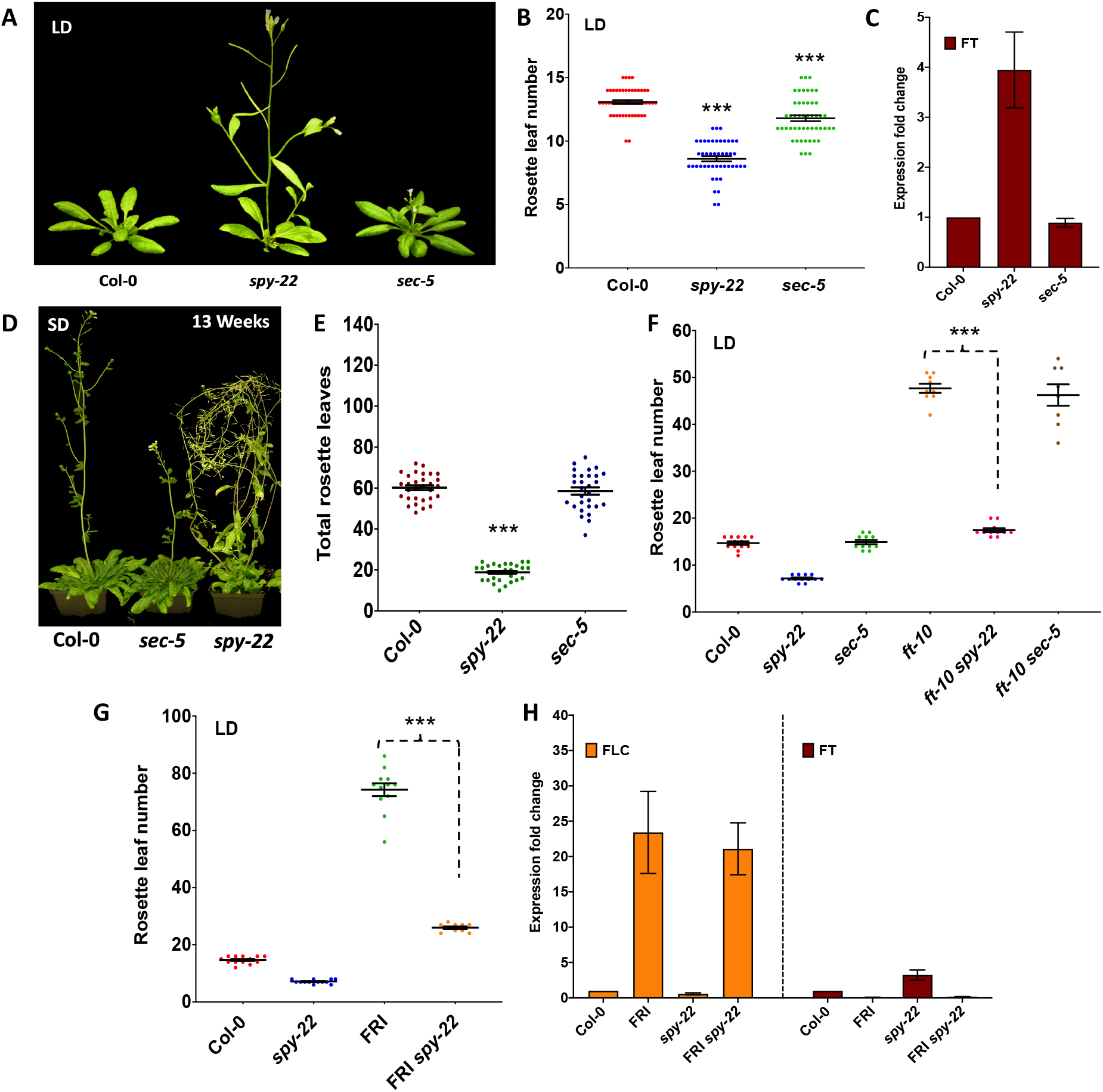
SPY suppresses flowering in long and short photoperiods. **(A)** (Representative pictures of wildtype Col-0, *spy-22* and *sec-5* grown in LD conditions, and **(B)** total rosette leaf numbers of these lines. (**C**) Relative expression levels of *FT* in 10-day old seedlings of *spy-22* and *sec-5* grown in LD conditions. (**D**) Representative pictures of wildtype Col-0, *spy-22* and *sec-5* grown in SD conditions, and **(E)** total rosette leaf numbers of these lines. (**F**) Total rosette leaf numbers of wildtype Col-0, *spy-22, sec-5, ft-10, ft-10 spy-22* and *ft-10 sec-5* grown in LD conditions. **(G)** Total rosette leaf numbers of wildtype Col-0, *spy-22*, Col-0 FRI and FRI *spy-22* grown in LD conditions, and **(H)** relative expression levels of *FLC* and *FT* in 10-day old seedlings of Col-0, *spy-22*, Col-0 FRI and FRI *spy-22* grown in LD conditions. For expression analysis, an average of three biological repeats ± SEM is shown, n > 20. For comparison of rosette leaf numbers, one-way ANOVA with Tukey’s multiple comparison was done, n is given in Table 1 (*** p ≤ 0.001).

### Mutants in SPY show accelerated transition from juvenile to adult phase

Bypassing of the vernalization requirement of Col-0 FRI has previously been shown in lines expressing an miRNA-resistant stabilized version of SQUAMOSA PROMOTOR-BINDING PROTEIN-LIKE 3 (rSPL3) that lacks an miRNA-binding site [9]. SPL-transcription factors regulate the juvenile to adult vegetative phase transition as well as flowering time; thus, we analyzed also the juvenile to adult vegetative phase change in O-glycosylation mutants. This transition is morphologically marked by the formation of trichomes on the abaxial side of leaves and changes in leaf morphology, an increase in the ratio of length to width of leaf blades, and the onset of serration [48] [49]. Based upon appearance of trichomes on the abaxial side of rosette leaves, we observed that *spy-22* plants consistently transitioned early from the juvenile to the adult stage at 3.5 ± 0.6 juvenile leaves (JL) compared to 5.8 ± 0.6 JL in Col-0, while *sec-5* (5.9 ± 0.7 JL) did not show significant differences compared to the wild-type (Figure 2A, Table 2). In order to dissect the juvenile to adult phase transition from floral transition, we included *ft-10-spy-22* in this analysis. We found that *ft-10* had a delayed juvenile-to-adult transition, with 9.2 ± 1.0 JL, which had been shown before [50]. On the other hand, *ft-10 spy-22* was similar to *spy-22* with 3.5 ± 0.6 JL, even though *ft-10 spy-22* (17.4 ± 1.3 TRL) showed an extended adult vegetative phase and was flowering considerably later than *spy-22* (7.1 ± 0.6 TRL) (Figure 2B, Table 2). This suggests that phase transition is regulated independently of daylength-dependent flowering time in *spy-22*. Juvenile-to-adult phase transition is regulated by an internal developmental program, orchestrated by a balance of counteracting miR156 and miR172 [7, 51]. Thus, we analyzed transcript levels of primary miRNA156a, miRNA156b and miRNA156c in 5-day old seedlings, but did not find differences between the wild-type and *spy-22* (Figure 2C).

**Table 2:**
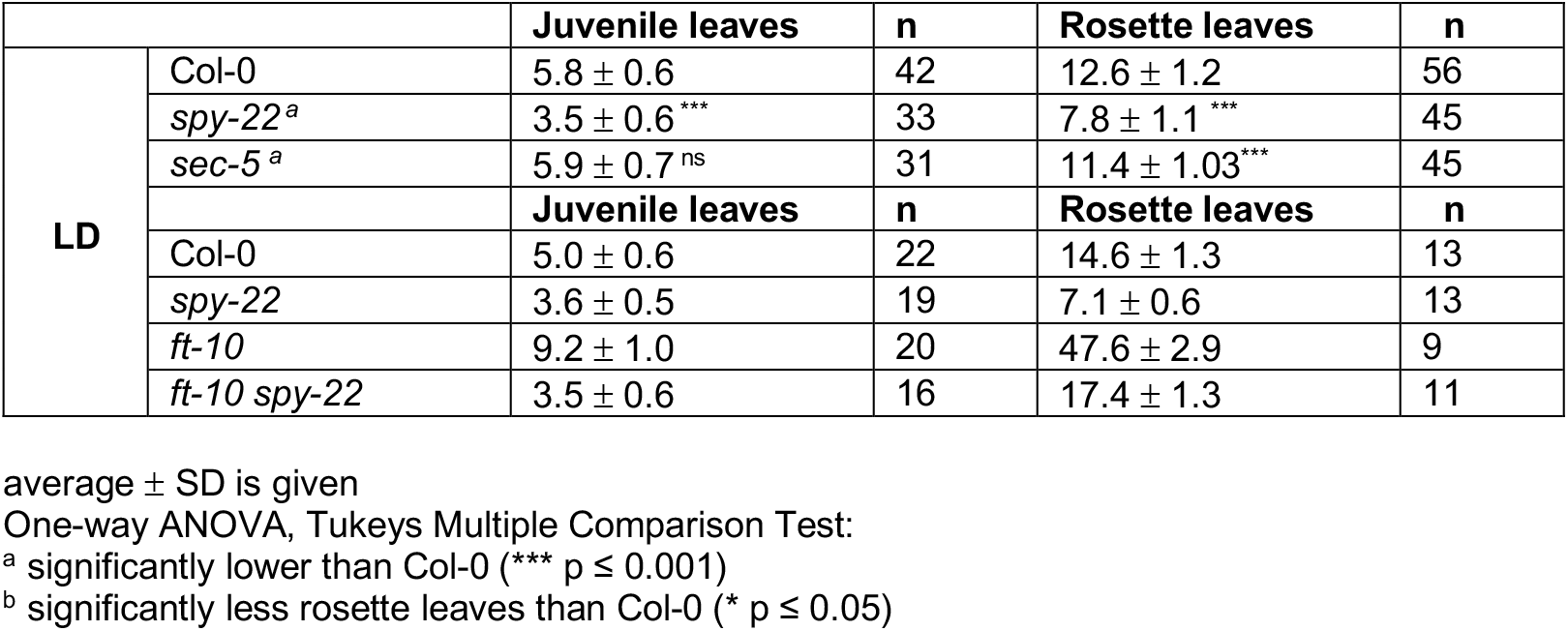
Rosette leaf numbers for graphs shown in Figure 2.

**Figure 2.**
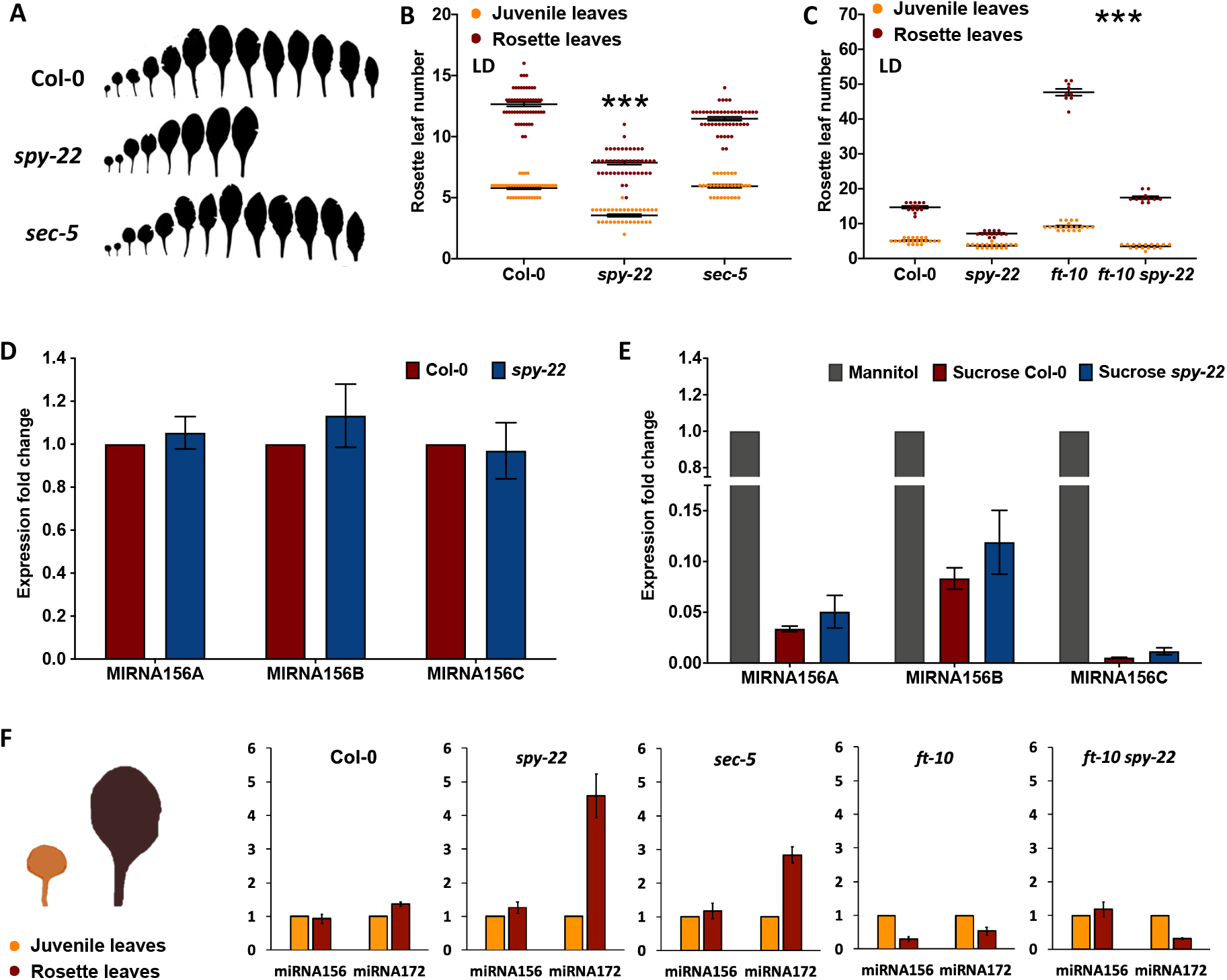
SPY regulates juvenile to adult phase transitions. (**A**) Leaf scans (left panel) and juvenile and total leaf numbers (right panel) of wildtype Col-0, *spy-22* and *sec-5* grown in LD conditions. One-way ANOVA with Tukey’s multiple comparison was done, n is given in Table 2. (*** p ≤ 0.001, ns: not significant). (**B**) Juvenile and total leaf numbers of wildtype Col-0, *spy-22, ft-10* and *ft-10 spy-22* grown in LD conditions. (**C**) Relative expression levels of primary *miR156a, -b* and *-c* in Col-0 and *spy-22* in 5-day old seedlings, an average of three biological repeats +/-SEM is shown, n > 20. **(D)** Relative expression levels of primary *miR156a, -b*, and *-c* in Col-0 and *spy-22* in 5-day old seedlings treated with 50 mM sucrose for 24 h. An average of two biological repeats is shown, n > 20. **(E)** Relative expression of mature *miR156* and *miR172* in juvenile and adult leaves of Col-0, *spy-22, sec-5, ft-10* and *ft-10 spy-22*. Representative results of one out of three biological repeats (average ± SD) are shown.

Levels of miR156 are, among other pathways, regulated by the accumulation of sugars in the leaves of growing plants [16, 17]. Cellular nutrient availability, especially increased sugar levels, enhance global levels of O-glycosylation in animals [52-55]. To test if O-glycosylation might be involved in regulation of miR156 abundance in response to sugar levels in plants, we treated 5-day old Col-0, *spy-22* and *sec-5* seedlings with 50 mM sucrose by transferring them to liquid media with and without added sucrose for 24 h. While we could reproduce the results of a strong decrease in miR156a, miR156b and miR156c in response to sucrose treatment as shown before [16, 17], we did not see any significant differences between *spy-22* and the wild-type after the treatment (Figure 2D).

We then compared expression of mature miR156 and miR172 in juvenile and adult leaves of soil-grown plants. Leaves were harvested for all lines at the timepoint when the wild-type plants started to bolt. We observed balanced levels of miR156 and miR172 between juvenile and adult leaves in Col-0. *spy-22*, which was already flowering, showed a strong shift in the balance towards miR172, with a more than 4-fold increase in adult leaves compared to the juvenile leaves (Figure 2E). *sec-5* showed the same tendency, but at a lower level, while *ft-10* and *ft-10 spy-22*, which were not flowering yet when the samples were taken, showed lower levels of miR172 in adult compared to juvenile leaves (Figure 2E). This observation correlates with the flowering time of these lines (Figure 1D and 2B, Table 1) and their developmental stage at the timepoint of sampling, with miR172 levels being increased in the lines that were already in the reproductive phase and decreased in the lines remaining in the vegetative stage. We did not observe a change in the ratio of miR156 in juvenile leaves compared to adult leaves in *spy-22* or *sec-5* in comparison to the wild-type (Figure 2E). We repeatedly observed lower levels of miR156 in adult leaves of *ft-10*, but not in *ft-10 spy-22*, again correlating with the fact that *ft-10* is the line with the longest vegetative phase in this dataset and was far from transition at the timepoint of sampling.

Taken together, we did not see a change in levels of miR156 in *spy-22* or *sec-5*, but we observed an increase in miR172 levels in *spy-22* and sec-5 as a consequence of accelerated floral transition. Our results suggest that there is no involvement of O-fucosylation in regulating the levels or balance of miR156 and miR172 during developmental phase transitions.

### SPY regulates phase transitions via SPL transcription factors independently of miR156

To further test for a potential interaction between miR156 and O-glycosylation in the regulation of developmental transitions we generated *35S::MIRNA156a* lines. Overexpression of *miRNA156a* in Col-0 led to a strong delay of the juvenile-to-adult phase transition and extremely delayed flowering, as shown before [7, 9] (Figure 3A-D). We then crossed *35S::MIRNA156a* Col-0-with *spy-22* and *sec-5*, and the offspring of these crosses displayed a strong repression of the miR156 overexpression phenotype. While *35S::MIRNA156a* Col-0 formed 51.8 ± 6.8 leaves, *35S::MIRNA156a spy-22* was flowering at 9.2 ± 1.2 TRL and *35S::MIRNA156a sec-5* at 11.3 ±1.2 TRL (Figure 3 A - C). A comparison of expression levels of miR156 in juvenile leaves *35S::MIRNA156a spy-22* and *35S::MIRNA156a sec-5* with the respective parent lines *spy-22, sec-5, 35S::MIRNA156a* Col-0 and the wildtype Col-0 showed very high levels of mature miR156 in the crossed lines, with miR156 levels at the same level in *35S::MIRNA156a* Col-0 and *35S::MIRNA156a spy-22*, and weaker overexpression in *35S::MIRNA156a sec-5* (Figure 3D). These data show that the effect of overexpressing miR156 is strongly suppressed in both *spy-22* and *sec-5* and that the early flowering of these mutants is independent of miR156 levels.

**Figure 3.**
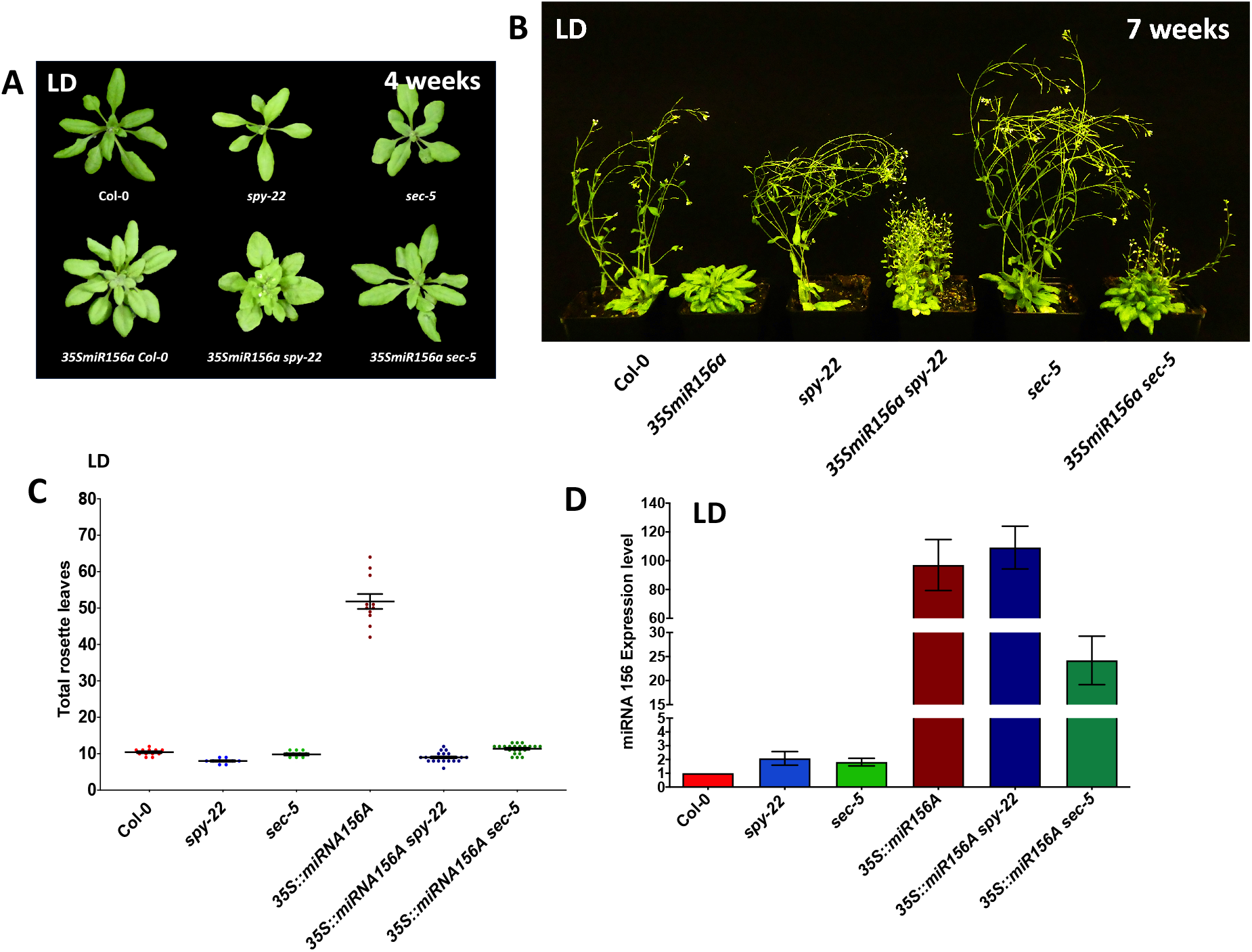
Overexpression phenotypes of miR156 are suppressed in *spy-22* and *sec-5*. **(A, B)** Representative pictures of 4-week-old (A) and 7-week-old (B) plants of wildtype Col-0, *spy-22* and *sec-*5 crossed with *35S::miRNA156a*. **(C)** Total rosette leaves of *35S::miRNA156a* Col-0, *35S::miRNA156a spy-22* and *35S::miRNA156a sec-5*, n is given in Table 3. **(D)** Relative expression levels of mature *miR156* in juvenile leaves of wildtype Col-0, *spy-22, sec-5, 35S::miRNA156a* Col-0, *35S::miRNA156a spy-22* and *35S::miRNA156a sec-5*. One biological repeat (average ± SD) is shown.

miR156 targets the family of SPL-transcription factors, which induce juvenile and adult phase transitions and floral transition [4, 7, 8, 51]. We therefore generated crosses of the O-glycosylation mutants with *spl9-4 spl15-1*, a line displaying delayed phase change, to test if SPLs are necessary for early developmental transitions of *spy-22*. We chose this line, as it has been characterized to show the strongest effect in delaying phase transitions among combinations of *SPL*-double mutants previously described [8], and both SPL9 as well as SPL15 directly interact with the DELLA protein RGA [11, 12], which is itself a target of SPY and SEC [38, 39]. We included *SPL9::rSPL9:GFP* [9] in our analysis, a line carrying an miR resistant version of SPL9 that consequently displays very early phase transitions. The early juvenile-to-adult transition of *spy-22* (3.5 ± 0.6 JL) was partly suppressed in the *spy spl9/15* triple cross (5.4 ± 0.7 JL), and flowering time was also affected (7.8 ± 1.1 TRL in *spy-22*, and 10.2 ± 1.5 TRL in *spy spl9/15*, Figure 4A-B, Table 4). In *sec spl9/15*, the number of juvenile leaves was slightly increased compared to *sec-5* (5.9 ± 0.7 JL in *sec-5* and 6.8 ± 0.7 JL in *sec spl9/15*), and *spl9/15* also delayed flowering of *sec-5* in the triple mutant (11.4 ± 1.03 TRL in *sec-5* and 16.3 ± 1.5 TRL in *sec spl9/15*) indicating that SPY and SEC might have common functions in this pathway (Figure 4A-B, Table 4). We did not see a full suppression of the early flowering of *spy-22* in *spy spl9/15* to the level of *spl9-4/15-1*. SPLs are encoded by a gene family with partially overlapping functions [8, 15], and potentially higher order *SPL*-mutants could lead to a stronger effect. Also, the effect of SPY on the photoperiod pathway and *FT* expression function independently and in parallel to the aging pathway, accelerating flowering time in parallel to SPLs.

**Table 3:** Rosette leaf numbers for graphs shown in Figure 3.

| | Rosette leaves (average ± SD) | n |
| --- | --- | --- |
| Col-0 | 10 ± 0.9 | 12 |
| spy-22 | 8 ± 0.6 | 10 |
| sec-5 | 9.8 ± 0.9 | 11 |
| 35S::MIR156a Col-0 | 51.8 ± 6.8 | 11 |
| 35S::MIR156a spy-22 | 9 ± 1.2 | 24 |
| 35S::MIR156a sec-5 | 11.3 ± 1.2 | 22 |

average ± SD is given

**Table 4:** Rosette leaf numbers for graphs shown in Figure 4.

|  |  | Juvenile leaves | n | Rosette leaves | n |
| --- | --- | --- | --- | --- | --- |
| LD | Col-0 | 5.8 ± 0.6 | 42 | 12.6 ± 1.2 | 56 |
|  | *spy-22*<sup>d</sup> | 3.5 ± 0.6 | 33 | 7.8 ± 1.1 *** | 45 |
|  | *sec-5*<sup>d</sup> | 5.9 ± 0.7<sup>ns</sup> | 31 | 11.4 ± 1.03*** | 45 |
|  | *spl9-4/15-1*<sup>e</sup> | 8.8 ± 0.6 | 37 | 19.6 ± 2.2 | 49 |
|  | *spy-22 spl9-4/15-1*<sup>a b</sup> | 5.4 ± 0.7 | 42 | 10.2 ± 1.5 | 54 |
|  | *sec-5 spl9-4/15-1*<sup>a c</sup> | 6.8 ± 0.7 | 46 | 16.3 ± 1.5 | 60 |
|  | SPL9::rSPL9:GFP | 1.3 ± 0.5 | 37 | 5.4 ± 1.1 | 50 |

average ± SD is given
One-way ANOVA, Tukeys Multiple Comparison Test:
<sup>b</sup> significantly less juvenile and rosette leaves than *spl9-4/15-1* (\*\*\*) $p \leq 0.001$
<sup>c</sup> significantly less juvenile leaves than *spl9-4/15-1* (\*\*) $p \leq 0.01$
<sup>d</sup> significantly less rosette leaves than Col-0 (\*\*\*) $p \leq 0.001$
<sup>e</sup> significantly more rosette leaves than Col-0 (\*\*\*) $p \leq 0.001$

**Figure 4:**
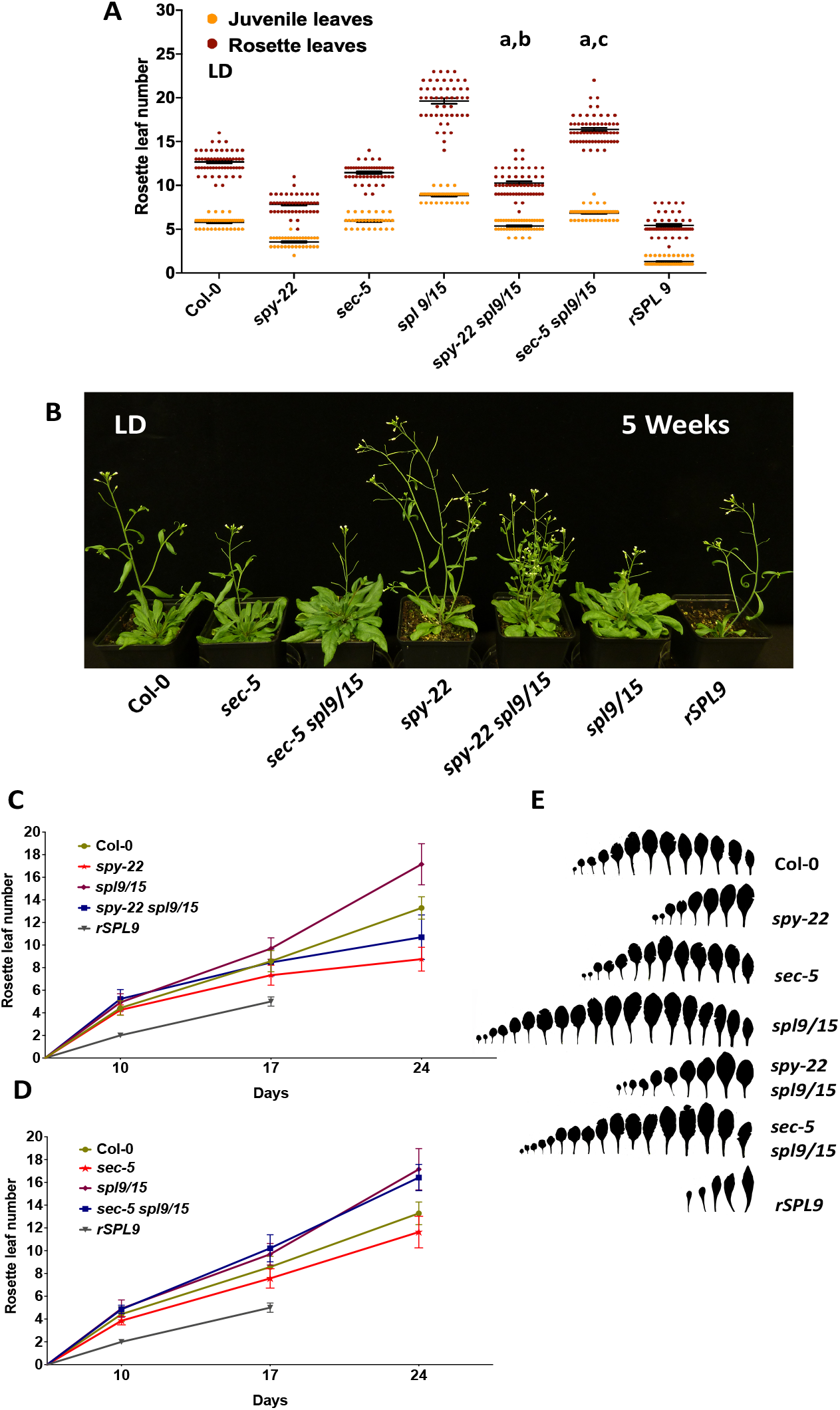
O-glycosylation represses phase transitions partly via SPL9 and SPL15. **(A)** Juvenile and total rosette leaf numbers of plant lines indicated in the graph, grown in LD conditions. Averages ± SEM are shown. For comparison of rosette leaf numbers, one-way ANOVA with Tukey’s multiple comparison was done, n is given in Table 4 (^a^ significantly more juvenile and rosette leaves than *spy-22* or *sec-5* (*** p ≤ 0.001), ^b^ significantly fewer juvenile and rosette leaves than *spl9-4/15-1* (*** p ≤ 0.001) ^c^ significantly fewer juvenile leaves than *spl9-4/15-1* (** p ≤ 0.01). Full statistical comparison is given in Table 4. **(B)** Representative pictures of 6-week-old plants with leaf numbers shown in (A). **(C**,**D)** Rosette leaf numbers over time of plants quantified in (A), averages ± SD are given. **(E)** Leaf scans of total rosette leaves of the indicated lines.

We also observed additive effects in shoot architecture in the triple crosses (Figure 4B). While *spy spl9/15* showed a strong loss of apical dominance compared to the wildtype, we observed the opposite in *sec spl9/15*, with a strong increase in apical dominance. While we did not follow up or quantify these observations yet, this might be an indication that SPY and SEC function redundantly as well as independently in different pathways.

We also observed the plastochron length by quantifying the number of leaves formed in regular intervals in parallel to determining the absolute leaf number (Figure 4C-D). *SPL9::rSPL9:GFP* was previously described as having a long plastochron, producing fewer leaves per day than wildtype, while *spl9/15* has a short plastochron [9], which we also observed. In this analysis, both *spy-22* and *sec-5* show a slightly longer plastochron compared to the wildtype, which was suppressed in both triple crosses with *spl9/15* (Figure 4C-D).

Taken together, our results suggest that SPY and SEC both negatively regulate phase change and flowering independently of miR156, potentially in interaction with SPLs.

### SPY and SPL9/15 are involved leaf size determination

While quantifying flowering time of the distinct mutants, we noticed additional shoot growth phenotypes that were not observed in the parent lines. In order to analyze these in more detail and dissect putative additive effects, we determined individual leaf areas as well as rosette areas and rosette growth rates (Figure 5). At 25 days after sowing (DAS), *spy-22* showed strongly decreased leaf areas, while *sec-5* had only slightly smaller leaves (Figure 5A, B). The *spl9/15* mutant had produced more leaves than the wildtype at this timepoint, correlating to the shorter plastochron (Figure 4C and 4D). Additionally, *spl9/15* showed a strong decrease in the areas of the first 7 leaves, while the leaf areas from leaf 9 onwards were increased compared to the wild-type Col-0, suggesting a developmental shift. This shift was also observed in the triple crosses with *spy-22* and *sec-5*. Additionally, we observed a clear additive effect for the decrease in leaf size, suggesting that O-glycosylation and SPL9 and SPL15 regulate leaf size independently from each other, but the developmental shift seen in *spl9/15* is epistatic in the triple crosses (Figure 5B). *SPL9::rSPL9:GFP* showed the highest increase in leaf size (with leaf 1 and 2 showing an increase in leaf area of 121% compared to Col-0) and a developmental shift compared to the wild-type in the opposite direction of *spl9/15*. Also in contrary to *spl9/15*, the number of leaves produced in *SPL9::rSPL9:GFP* at 25 DAS was strongly reduced to 3.9 ± 0.3 leaves, compared to 13.0 ± 1.0 leaves in the wildtype, similar to the increased plastochron observed in (Figure 4C-D), and [9]. Accordingly, the rosette area calculated as the sum of the individual leaf areas (mm^2^) was drastically reduced in *SPL9::rSPL9:GFP* (Figure 5A,C). At 25 DAS, *spy-22* and *spy-22 spl9/15* also showed a strong reduction in rosette area, while rosettes of *sec-5* were only slightly reduced (Figure 5A-B). The rosette areas of *spl9/15* and *sec spl9/15* were not significantly affected (Figure 5 A-C).

**Figure 5.**
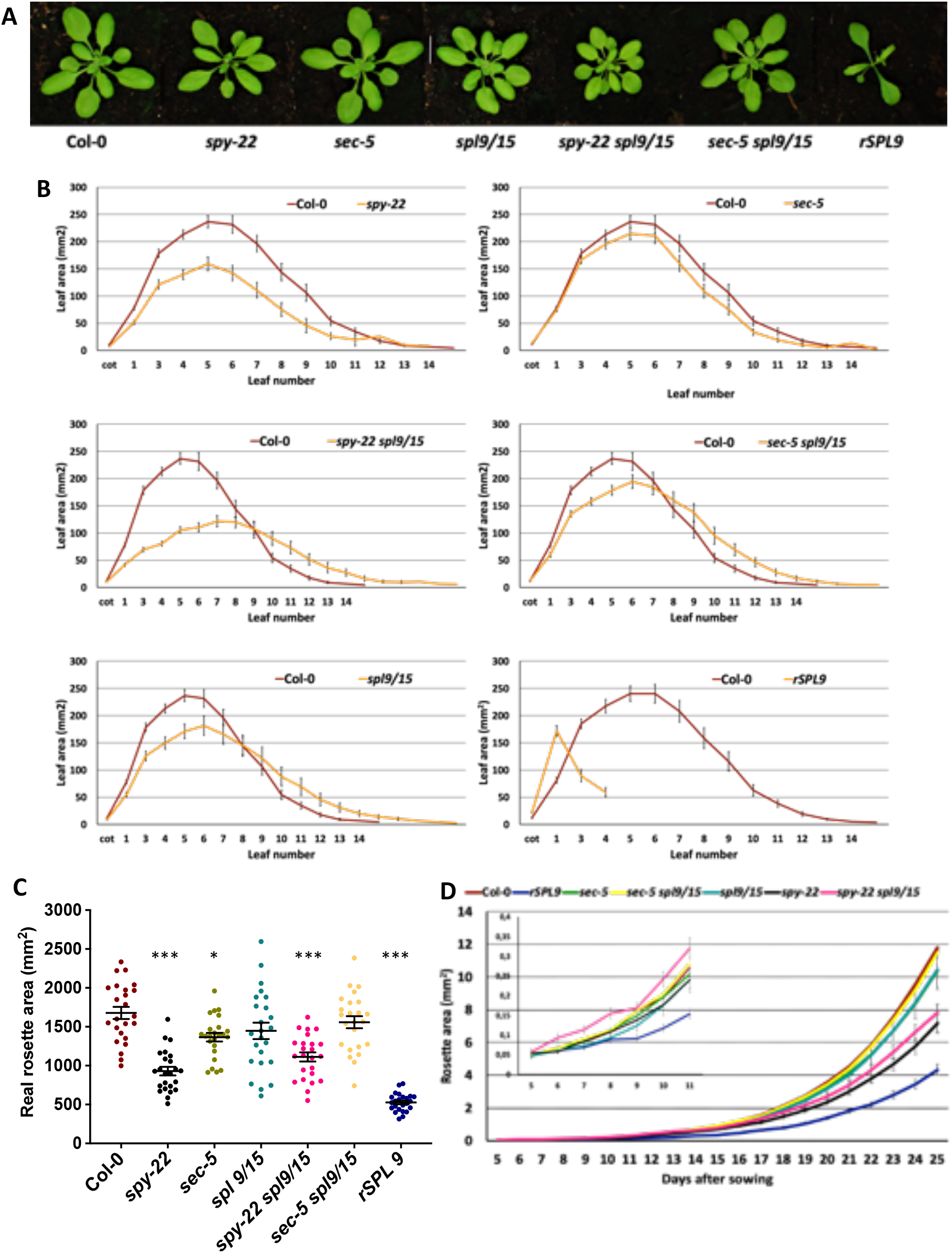
SPY and SPL9/SPL15 regulate leaf growth. (**A)** Representative pictures of 25-day-old rosettes of plants used for leaf and rosette area measurements. **(B)** Leaf areas of lines indicated in the graph. Averages ± SEM of three biological repeats using 8 plants per line for each repeat are shown. **(C)** Real rosette area, calculated from the leaf area measurements depicted in (B). For statistical analysis, one-way ANOVA with Tukey’s multiple comparison was done, significant differences to the wildtype Col-0 are shown (*** p ≤ 0.001, * p ≤ 0.05). **(D)** growth rates of rosette areas (mm^2^) of the indicated lines.

In a time course analysis, rosette growth correlated to the plastochron length in the analyzed lines (Figure 4C and D), with *SPL9::rSPL9:GFP* showing the slowest growth (Figure 5C-D). This is mainly resulting from the low number of leaves, as the individual leaf size for the first developing leaves was increased strongly in this line compared to Col-0 (Figure 5B). In *spl9/15*, the decrease in leaf size (Figure 5B) was partly compensated by the higher number of leaves due to shorter plastochron (Figure 4C and D). A positive correlation of plastochron and leaf size has also previously been shown in [56]. *spy-22* and *spy spl9/15* showed a considerably slower rosette growth compared to the wildtype, due to decreased leaf size. *sec-5* and *sec spl9/15* showed rosette growth rates comparable to the wildtype (Figure 5D).

To determine the cellular effect of the differences in leaf areas observed in O-glycosylation and *spl* mutants, we determined epidermal cell size and number in the first leaf pair (L1/2) of all lines. Leaves were harvested at 25 DAS, a timepoint when wild-type Col-0 leaves are considered mature. The leaves were cleared and microscopic drawings of the cells were used to determine pavement-, guard- and total cell number and individual pavement cell area (Figure 6A). Total cell number (guard and pavement cells) was strongly increased in *SPL9::rSPL9:GFP* (Figure 6B, Fig. S2A-B), but there was no change in pavement cell area (Figure 6C), suggesting that the increase in leaf area in *rSPL9* mutants is due to increased cell division. In all other lines, guard- and pavement cell numbers were not significantly changed compared to the wildtype (Figure 6B, Fig. 6SA-B). When measuring individual pavement cell areas (PCA), we found a decrease in *spy-22* (−34%) and *spy spl9/15* (−47%) compared to the wild-type, with a clear additive effect. PCA was also decreased in *spl9/15* (−30%) and *sec spl9/16* (−23%) mutants, though not statistically significant. The measurements go in line with the decrease in leaf and rosette area described above, suggesting that the leaf area decrease in *spy-22, spl9/15* and *spy spl9/15* results from additive defects in cell expansion (Figure 5B). However, we cannot exclude that the observed effects are not due to slower leaf growth rates on cellular level in these lines. Analysis of mature leaves in *spy-22* and *spy spl9/15*, taking samples at a later timepoint, could further address this question. Overall, however, our data suggest that cell division is not affected in these mutants. Given the increased cell division in *SPL9::rSPL9:GFP* (Figure 6A-C), we cannot exclude that redundancy among SPLs masks potential effects on cell division in *spl9/15*. Analysis of higher order mutants would address this question.

**Figure 6.**
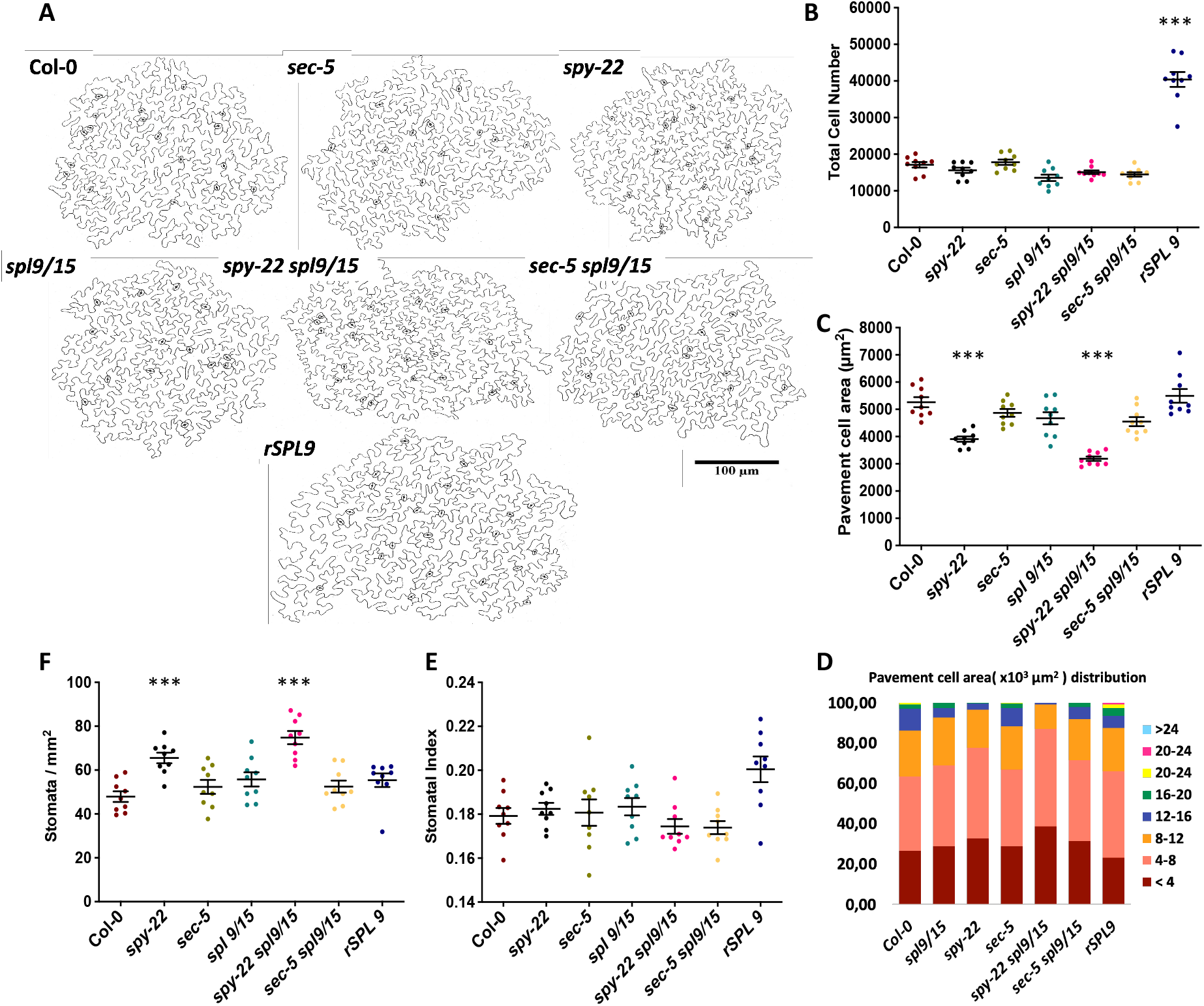
Cellular analysis of *spy-22, sec-5, spl9/15 triple mutants and SPL9::rSPL9:GFP*. **(A)** Representative drawings of cell margins of the indicated lines, as used for the analysis of cell numbers, areas, circularity, SI and SD. **(B)** Total epidermal cell number and **(C)** pavement cell area of lines indicated in the graph. **(D)** Distribution of pavement cell area of the indicated lines. **(E)** Stomatal Index (SI) and **(F)** stomata density (SD) of the indicated lines. Averages ± SEM of three biological repeats using 3 plants per line for each repeat are shown (n = 9). For statistical analysis, one-way ANOVA with Tukey’s multiple comparison was done, significant differences to the wildtype Col-0 are shown (*** p ≤ 0.001, * p ≤ 0.05).

The distribution of cell sizes indicates fewer cells in the category of the biggest cells in *spy-22*, and especially in *spy spl9/15* (Figure 6D), correlating with the results of the decreases in leaf area (Figure 5B). The number of cells in the smallest category is increased in these lines, suggesting a developmental shift at cellular level. Again, however, we cannot exclude that this is potentially due to cells not yet having reached maturity in these lines.

We then also determined stomatal index (SI) and stomata density (SD), corresponding to the proportion of stomata per pavement cells and the density of the stomata, respectively. For SI, we only observed a slight increase in *SPL9::rSPL9:GFP*, suggesting that more stomata are initiated, while all the other lines did not show significant differences compared with wildtype (Figure 6F). On the other hand, SD was significantly increased in *spy-22* and *spy spl9/15* (Figure 6G), likely to result from the decrease in average pavement cell area in these lines at the timepoint used for phenotypic analysis (Figure 6C).

Overall, our data suggest that SPL9 positively regulates cell division, while we did not see an effect on cell division in the other analyzed lines. We also observed a reduction in pavement cell area in *spy-22* and *spy spl9/15*, either due to defects in cell expansion or delayed growth rates.

### SPY O-fucosylates SPL transcription factors

Several SPLs were recently shown to be O-GlcNAc modified, among them also SPL8, which does not carry a miR156 recognition site [40]. We therefore tested if SPL15, which is important for the control of flowering time, directly interacts with SPY and can be O-fucosylated. *35S::FLAG:SPY* was co-infiltrated with *35S::SPL15:HA* and *35S::SPL8:HA* as a positive control, respectively, in *N. benthamiana* leaves for co-immunoprecipitation. When analyzing protein expression after infiltration of *N. benthamiana* leaves, we could clearly see a shift of SPL8 and SPL15 when co-infiltrated with SPY compared to the independent infiltration, suggesting a modification of SPL8 and SPL15 by SPY (Figure 7A). Interestingly we also consistently observed a stabilization of SPL8 and SPL15 in the samples co-infiltrated with SPY compared to single infiltrations (Figure 7A, a representative example of several biological repeats is shown.) When precipitating Flag:SPY with an anti-Flag antibody, we could pull down SPL15, indicating protein-protein interaction (Figure 7B). These results indicate that SPY directly interacts with, and glycosylates SPL15. Together with our data from genetic analysis, this suggests that SPY inhibits SPL-activity by glycosylation.

**Figure 7.**
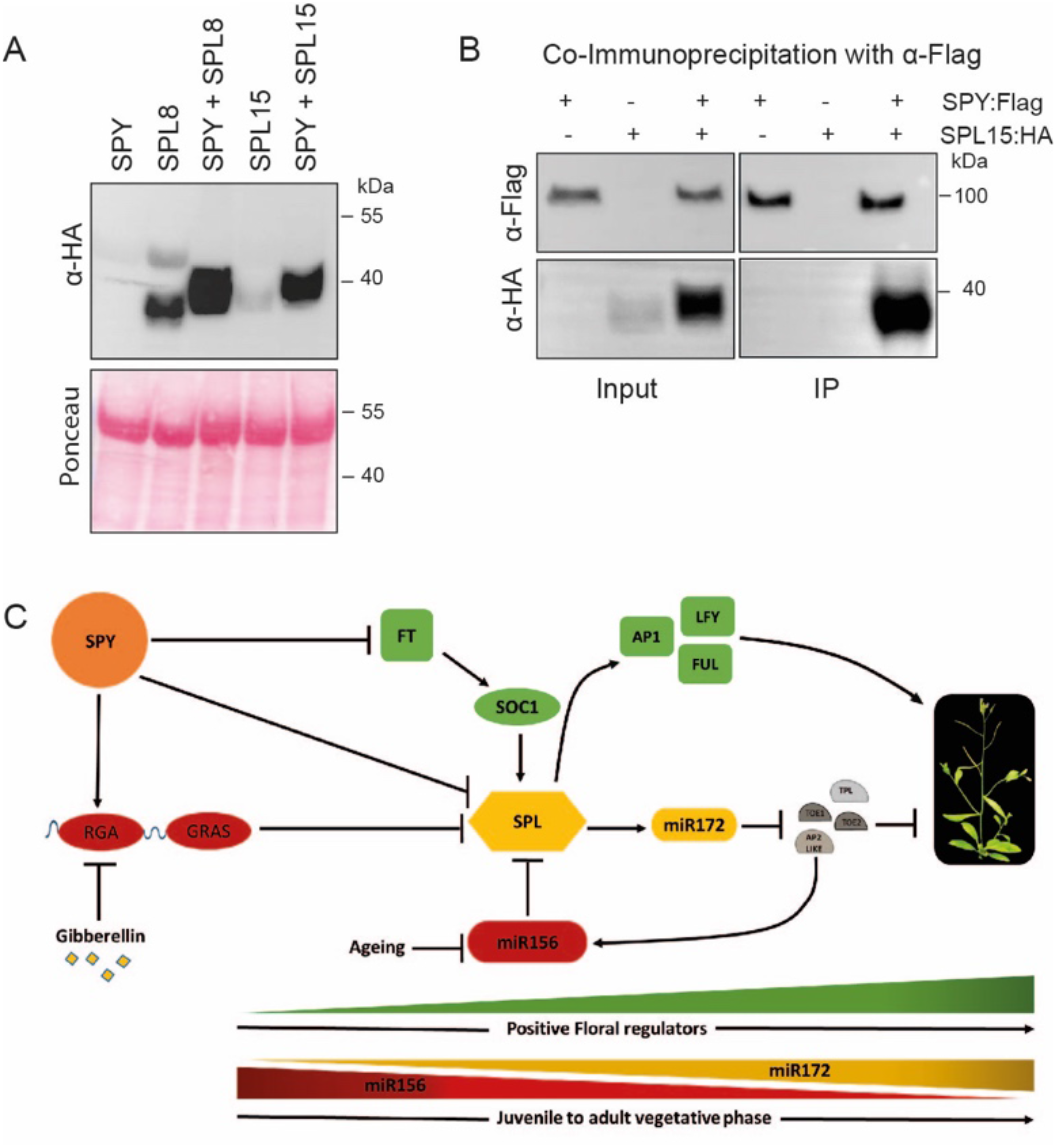
SPY interacts with SPLs. (**A)** Western Blot of protein extracts from transient expression in *N. benthamiana, 35S::SPY:Flag, 35S::SPL8:HA, 35S::SPL15:HA* single infiltrations, as well as *35S::SPY:Flag, 35S::SPL8:HA* and *35S::SPY:Flag, 35S::SPL15:HA* co-infiltration is shown. Protein extracts were blotted and probed with anti HA-antibody to visualize SPL8 and SPL15. **(B)** Samples after co-infiltration of transiently expressed *35S::SPY:Flag* and *35S::SPL15:HA* were used for immunoprecipitation of SPY using an anti-Flag antibody, co-immunoprecipitation of SPL15 is shown with an anti-HA antibody. **(C)** Current working model suggesting a role for O-glycosylation at several levels in the regulation of developmental transitions in *Arabidopsis thaliana*.

## DISCUSSION

A multitude of different post-translational modifications (PTMs) confer complexity to the regulation of protein function and stability. They are essential components of signalling pathways in the course of development, often integrating environmental changes or fine-tuning crosstalk between different regulatory pathways. O-glycosylation of cytosolic and nuclear proteins is the modification of a number of very diverse proteins with a single monosaccharide on serine or threonine residues – which is in contrast to N-glycosylation events that involve the formation of branched carbohydrate chains of varying composition in the secretory pathway [57]. Most organisms carry only one type of cytosolic O-glycosylation, with O-GlcNAcylation being the most common and best described example that is very well conserved among all kingdoms. Yeast is lacking an O-GlcNAc transferase, but uses instead O-mannose modification for the same molecular function. Many of the target proteins are however conserved between yeast and animal cells [58, 59], suggesting that the molecular function is more conserved than the type of monosaccharide involved. Plants are exceptional in that they use two different sugars attached to the same protein targets [39]. Currently the only other organism described to use fucosylation as well as GlcNAc modification is *Toxoplasma gondii*, but probably with distinct targets for the two different modifications [59].

In Arabidopsis, only few O-glycosylated targets have been characterized so far. O-glycosylation of the transcription factors TEOSINTE BRANCHED, CYCLOIDEA AND PCF 14 (TCP14) and TCP15 by SPY increases their stability during cytokinin signalling [60, 61]. SPY was also implicated in the integration of reactive oxygen species signalling during root development [62]. Further protein interactions between SPY and GIGANTEA [44], and SPY and SWI3C [63] have been identified. However, by far the best characterised O-glycosylated protein in plants is the DELLA protein RGA. The current model suggests that O-GlcNAc and O-fucose modification have opposite effects on DELLAs during GA signalling, with O-GlcNAcylation leading to a closed conformation of DELLAs, rendering them less active. On the other hand, fucosylation leads to an open conformation of RGA, facilitating the interaction with target transcription factors, thus increasing DELLA activity [38, 39]. However, a number of open questions remain, even in the context of GA signalling, such as the observation that the double knockout of SPY and SEC is embryonic lethal even using weaker alleles, while the single mutants don’t show drastic developmental phenotypes [64]. This suggests that there might be targets where both modifications have the same effect, or even both modifications are necessary at the same time, though not identified so far. A proteomic study using lectin weak affinity chromatography (LWAC) with glucosamine-binding wheat germ agglutinin (WGA) revealed O-GlcNAc modification of many different proteins, among them a number of transcriptional regulators, including SPLs [40]. SPL9 and SPL15 directly interact with RGA [11, 12, 32], and this interaction is likely to be affected by glycosylation of RGA as it was shown for other transcription factors such as BRASSINAZOLE-RESISTANT 1(BZR1), JASMONATE-ZIM-DOMAIN PROTEIN 1 (JAZ1), PHYTOCHROME INTERACTING FACTOR 3 and 4 (PIF3, PIF4) [38, 39]. Here, we show that SPL15 additionally directly interacts with and is modified by SPY, suggesting that there is an additional, potentially independent effect of glycosylation on SPL15 function. Moreover, we see an additive effect in *spy spl9/15* on leaf size as well as pavement cell size, indicating that SPLs might additionally be regulated by other factors in the determination of leaf size.

Our data indicate that the activity of SPLs is stabilized in both O-glycosylation mutants despite overexpression of *miR156a*, as the late flowering phenotype of *35S::miR156a* is almost completely suppressed in *spy-22 35S::miR156a* and *sec-5 35S::miR156a*. This is supported by the genetic analysis using *spl9/15* crosses with *spy-22* and *sec-5*, where the early flowering of *spy-22* and *sec-5* is partly suppressed, suggesting that functional SPLs are necessary for the accelerated flowering time in O-glycosylation mutants. It has previously been shown that miR156 overexpression lines are less sensitive to GA treatment in terms of flowering time, keeping their extreme late flowering phenotype even when sprayed with GA [9, 11]. *spy*-mutants are often described as showing constitutive gibberellin signalling, but in contrast to gibberellin treatment of miR156 overexpression lines, we see a strong suppression of late flowering in *spy-22 35S::MIR156a* mutants. Moreover, albeit SPY and SEC have opposite effects in gibberellin signalling [38, 39], *sec-5* as well as *spy-22* suppress the late flowering of *35S::MIR156a*, and to a lesser extent developmental transitions, suggesting that the effect of glycosylation on SPLs might be independent of GA signalling and DELLAs, as well as miR156, with a degree of redundancy between O-GlcNAc and O-fucose.

Phenotyping of leaf and rosette growth as well as cellular analysis of our lines suggest a role for both SPY and SPL9/SPL15 in leaf and cell size, with a clear additive effect. Again, we observed a potentially redundant function for SPY and SEC in this respect, although the phenotypes of *sec-5* were very subtle, and not statistically significant. While we did not see an effect of SPY on cell division, we observed a strong positive effect on cell division in *SPL9::rSPL9:GFP*. On the other hand, total cell number was not significantly decreased in *spl9/15*. This may be due to redundancy and potentially higher order *spl* mutants might show a defect in cell division. However, the results of the leaf and rosette growth analysis might reflect different plastochron length or rates of growth and/or development rather than organ size regulation in the analyzed lines. Sampling of leaves at a later timepoint, when the leaves of all lines are fully developed and have reached maturity, might reveal additional cellular effects in these mutants and lead to additional conclusions. It will be interesting to see in further studies, if (1) SPY indeed regulates cell expansion, but not cell division, via SPLs, or if (2) the differences we see in leaf size are a result of different developmental timing and leaf growth rates in *spy-22* and *spl9/15* mutants, or (3) if the effect of SPY on cell expansion is completely independent of SPLs, for instance by interaction with other targets.

Overall, we suggest a model, where O-glycosylation regulates developmental transitions on multiple levels (Figure 7C). In long photoperiods, SPY suppresses expression of *FT*, via interaction with GI [44]. Additionally, glycosylation of RGA regulates its interaction with transcription factors regulating flowering time, such as PIFs [35, 39, 65] and potentially also SPLs [11, 12, 32]. Our data add an additional level of regulation by direct glycosylation of SPLs, controlling them independently of miR156, and potentially also independently of gibberellins.

## MATERIAL AND METHODS

### Plant Material and growth conditions

*Arabidopsis thaliana* ecotype Col-0 of was used as wild type. Lines *spy-22* (SALK_090582), *sec-5* (SALK_034290), *spl9-4 spl15-1* (N67865, [14]), *ft-10* (GK-290E08, [66]), FRI SF-2 (N6209) and pSPL9::GFP-rSPL9 (N9954, [9]), all in Col-0 background, were obtained from the European Arabidopsis Stock Centre (NASC) (Scholl et al., 2000). Seeds were surface sterilized with 70% ethanol and transferred ½ Murashige and Skoog medium (2.15 g/L MS Salts, 0.25 g/L MES, pH 5.7, 1% agar). Seeds were stratified in the dark at 4°C for 48 hours. Based upon the experiment the seedlings were germinated and grown in either long day (LD, 16 hours light / 8 hours dark) or short day (SD, 8 hours light / 16 hours dark) conditions at 22°C. For studying phase change and flowering time, seedlings were transferred to soil at 5 days after germination. For all phenotyping experiments, transgenic lines and their wild-type controls were sown in pots filled with soil. After sowing, the seeds were stratified for 3 days at 4°C and grown at 21°C under a 16 h day/8 h night regime for 25 days. Experiments were performed with transgenic and wild type seeds harvested from plants that were grown in parallel.

### Flowering time and phase transition quantification

Flowering time was quantified by counting the total number of rosette leaves (TRL) produced. Phase transitions were studied by observing the appearance of abaxial trichomes at the lower side of rosette leaves. Rosette leaves without any abaxial trichomes were grouped as juvenile leaves (JL) and rosette leaves with abaxial trichomes were considered adult. Arabidopsis rosettes were harvested and individual leaves were taped to white paper and scanned for representations. Scanned pictures were edited to black and white using Paint 3D.

### RNA extraction and qPCR

For expression analysis, seedlings were grown on ½ MS plates. For studying the primary miRNA expression in response to sucrose treatment, sterilized seeds were germinated in 50 ml ½ MS liquid media with shaking at 140 rpm. At 5 DAG, seedlings were transferred to ½ MS media supplemented with and without 50 mM sucrose, and seedlings were harvested after 24 h. 200 mg of total plant material was used for total RNA extraction using SV Total Isolation System (Promega). 1µg of RNA was used for cDNA synthesis with the iScript™ cDNA Synthesis kit (Bio-Rad). GoTaq® qPCR Master Mix (Promega) was used for quantitative real-time PCR, primers are listed in Table 5 and data was analyzed with Bio-Rad CFX Manager and Microsoft Excel for relative quantification using the 2(-ΔΔC(T))-method [67]. For technical repeats, every sample was done in triplicates, representative results from one of at least two biological replicates are shown (as given in the figure legends). For studying mature miRNA expression, rosette leaves were harvested at the point when Col-0 plants were flowering. Harvested rosette leaves were separated into juvenile and adult leaves, ground in liquid nitrogen and total RNA from approximately 150 mg of ground leaf material was extracted with a ReliaPrep miRNA Cell and Tissue Miniprep System kit (Promega). A stem loop pulsed RT-PCR protocol [8, 68] was used to study the expression of mature miRNA using a Thermo Scientific RevertAid First Strand cDNA synthesis kit and stem-loop primers for miR156 and miR172 listed in Table 5, small nuclear RNA snoR101 was used as internal reference. Equal amounts of each RNA and 1 µM stem loop primer were incubated at 65°C for 5 minutes with 10mM dNTPs. RiboLock RNase Inhibitor and a 5X reaction buffer were added and amplification was done for 60 cycles with 30 s of 30°C, 30 s of 42°C and 50°C for 1 s respectively. PCR products were diluted 4x in nuclease free water, qPCR was done with the GoTaq® qPCR Master Mix (Promega). For technical repeats, every sample was done in triplicates. Following qPCR, the data was analysed with Bio-Rad CFX Manager and Microsoft Excel. Representative results from one of three biological replicates are shown.

**Table 5:** Distribution of pavement cell circularity and area.

| Distribution of pavement cell area |  |  |  |  |  |  |  |
| --- | --- | --- | --- | --- | --- | --- | --- |
| Cell area<br>(x10<sup>3</sup> μm<sup>2</sup>) | Col-0 | *spy-22* | *sec-5* | *spl9/15* | *spy-22*<br>*spl9/15* | *sec-5*<br>*spl9/15* | *rSPL9* |
| < 4 | 26,82 | 32,73 | 28,96 | 28,94 | 38,77 | 31,49 | 23,18 |
| 4-8 | 36,59 | 45,05 | 38,03 | 39,90 | 48,27 | 39,94 | 42,88 |
| 8-12 | 22,74 | 18,77 | 21,68 | 23,97 | 12,35 | 20,58 | 21,36 |
| 12-16 | 10,83 | 3,15 | 8,74 | 4,62 | 0,62 | 5,84 | 6,29 |
| 16-20 | 2,49 | 0,30 | 2,43 | 2,57 | 0,00 | 2,15 | 3,81 |
| 20-24 | 0,53 | 0,00 | 0,16 | 0,00 | 0,00 | 0,00 | 1,66 |
| >24 | 0,00 | 0,00 | 0,00 | 0,00 | 0,00 | 0,00 | 0,83 |
| Sum (%) | 100,00 | 100,00 | 100,00 | 100,00 | 100,00 | 100,00 | 100,00 |

**Table 6:**
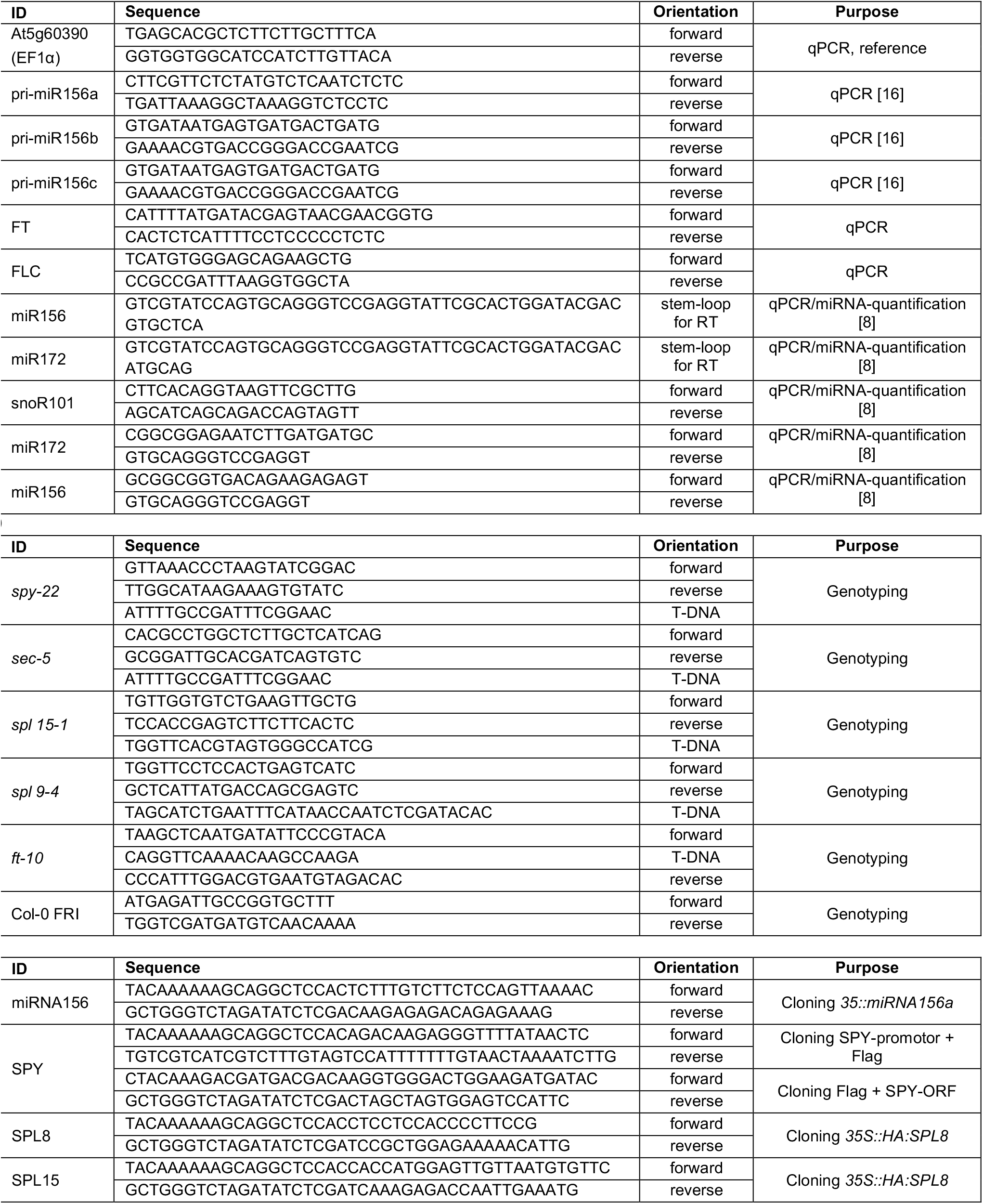
Oligonucleotides used in this study.

### Plasmid construction and generation of transgenic lines

*miR156a* overexpressing and *SPY::SPY:FLAG* constructs were generated by amplifying the genomic DNA from Col-0, and for transient overexpression, *SPY, SPL8* and *SPL15* were amplified from cDNA to generate the respective tagged constructs using Q5 high fidelity DNA polymerase (NEB). The primers (listed in Table 5) contained 5’-overhangs binding to the linearized, NcoI / XhoI-digested backbone of the cloning vector pENTR™ 4. PCR products were excised and purified from agarose gel using GeneJET Gel Extraction Kit (Thermo Fisher) and cloned into Gateway™ pENTR™ 4 by mixing the linearized vector backbone and PCR product in a 1:1 ratio using Gibson assembly (NEB), before transformation into DH10B electro-competent *E. coli* cells. Plasmids containing the gene of interest were extracted using GeneJET Plasmid Miniprep Kit (Thermo Fisher) and confirmed by sequencing.

Plant expression vectors were generated using the above created entry clones and destination vectors pK7WG2D (Karimi et al., 2002) for *35S::MIR156a*, pEarleyGate202 for *35S::FLAG:SPY*, pEarleyGate201 for *35S::HA:SPL8* or *35S::HA:SPL15* and pEarleyGate302 for *SPY::SPY:FLAG* (Earley et al., 2006). Recombination of the entry clones with the destination vectors was done using Gateway LR Clonase ll enzyme mix. Positive colonies with the plasmid of interest were selected for spectinomycin (150µg/mL) resistance for miRNA overexpressing pK7WG2D constructs, and kanamycin (50µg/mL) resistance for pEarleyGate201, pEarleyGate202 and pEarleyGate302 constructs, respectively on LB medium. Plasmids carrying the gene of interest were extracted from overnight bacterial culture using GeneJET Plasmid Miniprep Kit (Thermo Fisher) and confirmed by sequencing. Correct plasmids were transformed to *Agrobacterium tumefaciens* strain GV3101 (pMP90) before transformation of plants by floral dipping [69]. *35S:MIR156a* constructs were transformed to Col-0, then crossed to *spy-22* and *sec-5* backgrounds, and *SPY::SPY:FLAG* was introduced into *spy-22* background.

### Rosette growth and leaf series analysis

For the projected rosette measurements over time, pots containing the plants were positioned in a randomized way for 25 DAS on a platform, named MIRGIS, allowing fully automatic phenotyping. Pictures were acquired on a daily interval and image processing and analysis were performed using the ImageJ software (http://rsb.info.nih.gov/ij/). To construct the projected rosette area and rosette growth curves, the area measurements of similar transgenic lines plants were compiled. For the leaf series analysis, individual leaves (cotyledons and rosette leaves) of 25 day-old plants were dissected at the base of the petiole, oriented from old (cotyledons) to young on a thin layer of 1% w/v agar covering the bottom of a square plate (22.5 x 22.5 cm) and, if necessary, cut to flatten them. The leaf series were imaged and the individual leaf areas were measured with the ImageJ software (http://rsb.info.nih.gov/ij/). The real rosette area was defined as the sum of the individual plant leaves. The growth experiment was performed in three independent biological repeats containing 8 plants per experiment.

### Cellular analysis

Following the leaf series analysis at 25 DAS, the first leaf pairs (L1/2; 16 leaves) were cleared with 100% ethanol, stored in lactic acid and mounted on microscopic slides. The leaves were photographed with a dark-field binocular microscope and the leaf blade areas were determined with the ImageJ software. For each line, three leaves with an area closest to the median and the average were selected for detailed cellular analysis. Abaxial (lower) epidermal cells (∼150 cells) at the tip of the leaves were drawn with a binocular microscope (Leica, DMLB) fitted with a drawing tube and a differential interference contrast objective. The microscopic drawings were scanned for digitalization and guard, pavement and total cell numbers, as well as pavement cell area, stomatal density, stomatal index were calculated with the ImageJ software, as described in [19]. The cellular analysis experiment was performed in three independent biological repeats.

### Transient protein expression in *N. benthamiana*

For transient expression in *N. benthamiana, Agrobacterium tumefaciens* strain, GV3101 (pMP90) expressing *35S::FLAG:SPY* and *35S::HA:SPL8* or -*SPL15* constructs were cultivated overnight using rifampicin (50µg/mL), gentamycin (50µg/mL) and kanamycin (50µg/mL) selection. Bacterial cells were harvested, washed and resuspended in infiltration medium (500 mM MES, 20 mM Na_3_PO_4_, 100 µM acetosyringone, 50 mg/L D-glucose) to OD600 of 0.5. For co-infiltration, equal amounts of the respective cultures were mixed. After infiltration on the abaxial side of a leaf of a five-week old *N. benthamiana* plant, the plants were maintained in the growth chamber for 3 days before harvesting.

### Protein extraction and co-immunoprecipitation

For protein extraction, 1 g of agro-infiltrated leaves were frozen in liquid nitrogen, ground and taken up in 50 mM Tris-HCl pH 7.6, 150 mM NaCl, 1% Triton X-100, 1x plant protease inhibitor cocktail (Sigma) and 20 μM PUGNAc in the ratio 2:1. 100µL extract was stored for further analysis as input. 50 μL of anti-FLAG beads (Miltenyi Biotec) were added to the remaining plant extract and the samples were incubated on a rotor at 4°C for 30 minutes. A μMACS column was placed in the magnetic field of a μMACS separator and washed with above mentioned 200 μL extraction buffer. Plant extract containing the anti-FLAG beads was added to the column and 100 μL of the flow-through were collected and saved for further analysis. The column was rinsed 4 times with 200 μL TBS (50 mM Tris-HCl pH 7.5, 150 mM NaCl, 1x plant protease inhibitor cocktail (Sigma), 20 μM PuGNAc, 1.5 mM DTE) and once with 100 μL wash buffer (20 mM Tris-HCl pH 7.5). For elution, 20µL 0.1 M glycine, pH 2.3 was loaded onto to the column and incubated for 3-5 minutes. After adding 60µL 0.1 M glycine the first eluate was collected and neutralized with 20 µL 0.5 M Tris, pH 8.0. In the next step 20 μL of pre-heated 95°C hot Laemmli buffer (10% glycerol, 60 mM Tris-HCl pH 6.8, 2% SDS, 0.1% bromophenol blue, 5% β-mercaptoethanol) was loaded to the column and incubated for 5 minutes. Subsequently, 80 μL of pre-heated 95°C hot Laemmli buffer were added and the eluate was collected and used for SDS-PAGE analysis and Western blotting.

### Western Blotting and antibody dilutions

Input samples and eluates from the co-immunoprecipitation from both individual and co-infiltrated samples were subjected to SDS PAGE, and transferred to a PVDF membrane (Roth Roti®-PVDF, pore size 0.45 μm) using the wet transfer method in the Mini-Protean Tetra-System (Bio-Rad). The membranes were washed with PBST (137 mM NaCl, 2.7 mM KCl, 10 mM Na_2_HPO_4_, 1.8 mM KH_2_PO_4_, 0.1% (v/v) Tween 80) for 10 minutes, blocked with 3% milk for 1 hour at room temperature and probed with anti-FLAG M2 monoclonal antibody (mouse, Sigma F1804, 1:2000 in 3% milk/PBST), or blocked with 5% BSA for 1 hour at room temperature and probed with anti-HA antibody (rabbit, Cell Signaling Technology 3742S, 1:1000 in 5% BSA/PBST) respectively. After probing at 4°C overnight, the membranes were washed with PBST before incubation with the respective secondary antibodies, goat anti-mouse HRP (1:10000) (Dianova 115-035-164), and goat anti-rabbit HRP (1:20000, Agrisera A S09 602) for 1 hour at room temperature. After washing, Bio-Rad Clarity Western ECL substrate was used for chemiluminescence detection on a Fusion Solo S (Vilber).

### Data and statistical analysis

We used Excel for analysis of gene expression, and GraphPad Prism 5 and R (*ggplot*2, R Core Team, https://www.r-project.org/) for statistical analysis and generating graphs for quantification of rosette leaves. In the leaf quantification graphs, the mean is shown and error bars represent standard error, n is given in the respective tables. For statistical analysis, one-way ANOVA followed by Tukey’s multiple comparison test were done. For the leaf series analysis, statistical tests were performed with the mixed models and plm procedure using SAS Enterprise Guide 7 software as described previously in [70]. An F-test (p<0.05) was used to assess equality of variances. For each leaf, pairwise comparisons amongst the transgenic line or between the transgenic lines and the wild type control were performed using a Tukey-Kramer post-hoc analysis (p < 0.05). For the statistical analysis of the leaf blade area of L1/2 and the cellular data, ANOVA tests were performed in R (version 3.3.2) (https://www.r-project.org) followed by Tukey’s multiple comparison test (p < 0.05). For all statistical tests, the transgenic line was included as the fixed factor in the model. All experiments were performed in three biological replicates and the repeat factor was included as a random factor.

## ACKNOWLEDGEMENTS

We gratefully acknowledge Marie-Theres Hauser for sharing equipment, Lukas Grinninger for technical support and Julia König for supplying *N. benthamiana* plants. We thank Barbara Korbei, Vinod Kumar and Jürgen Kleine-Vehn for valuable input. This work was supported by the Austrian Academy of Sciences ÖAW (DOC-fellowship to K.V.M. and APART fellowship to D.L.) and the Austrian Science Fund FWF (P-29051). A.B. received funding from the Research Foundation Flanders (FWO) under the project number 3G038719.

## AUTHOR CONTRIBUTIONS

D.L. conceived the project, D.L., K.V.M designed experiments and wrote the manuscript. A.B. designed and performed phenotyping and cellular analysis, K.V.M. performed all other experiments with the help of N.N., C.F. and I.Z.

## DECLARATION OF INTERESTS

The authors declare no competing interests.

---

**Supplemental Figure 1.**
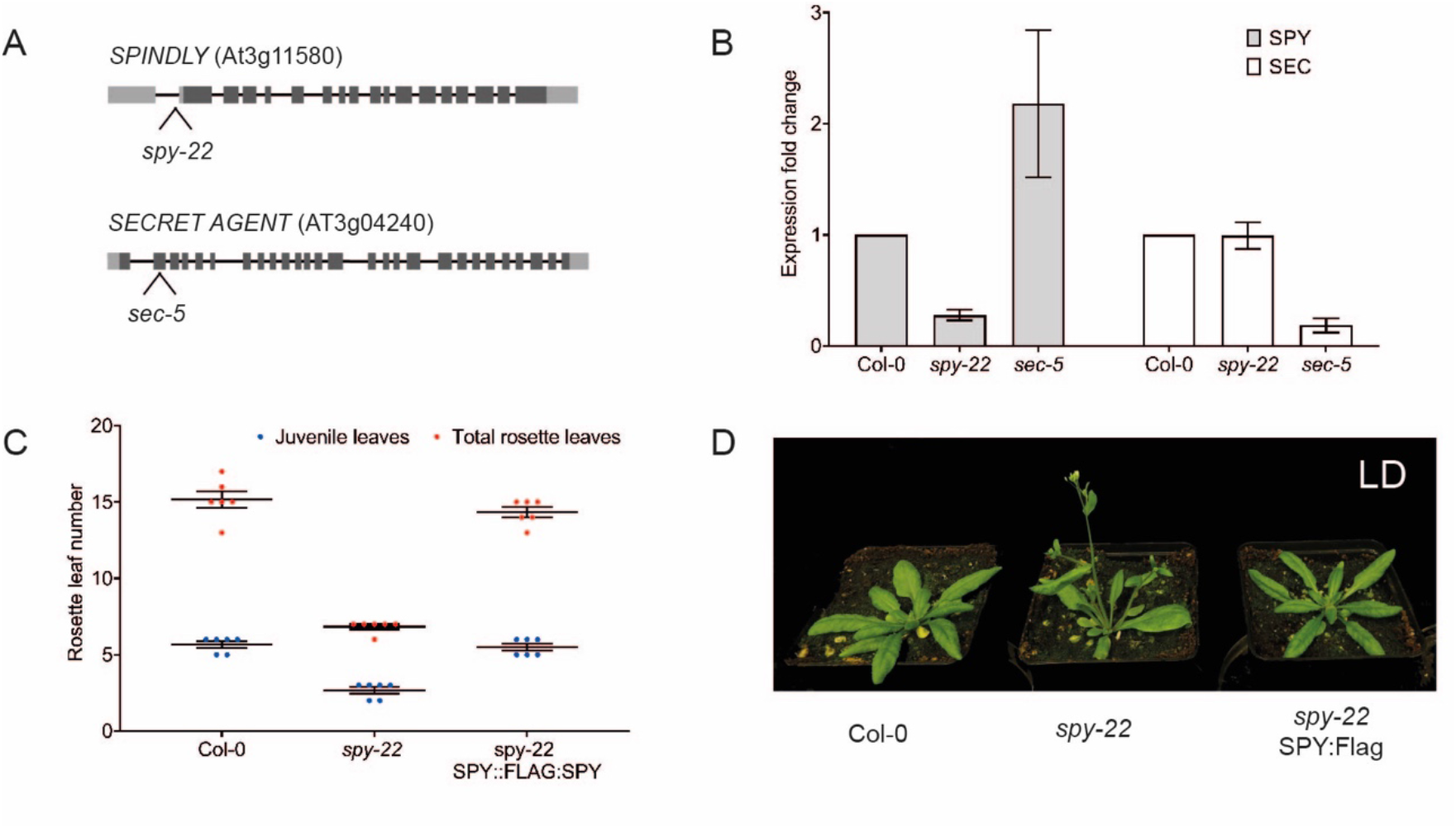
Description of mutant lines *spy-22* and *sec-5*. (**A**) Structure of *SPY* and *SEC*, 5’- and 3’-UTR are shown in light grey and exons in dark grey boxes, the lines represent introns. The position of the T-DNA insertion in *spy-22* (SALK_090582) and *sec-5* (SALK_034290) are given. (**B**) Relative expression levels of *SPY* and *SEC* in *spy-22* and *sec-5 seedlings*, an average of three biological repeats ± SEM is presented, n > 20. (**C**) Juvenile and total rosette leaf numbers of wildtype Col-0, *spy-22* and *spy-22 SPY::SPY:Flag* (*SPY:Flag*) grown in LD conditions. (**D**) Representative pictures of wildtype Col-0, *spy-22*, and *spy-22 SPY::SPY:Flag* (*SPY:Flag*) grown in LD conditions.

**Supplemental Figure 2.**
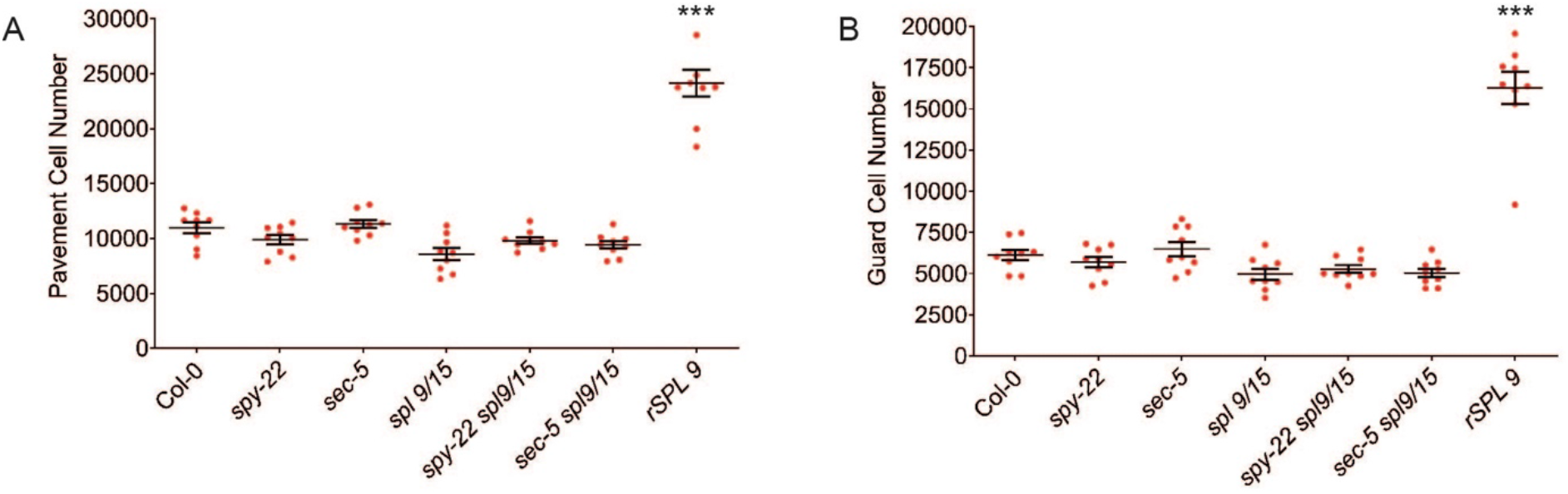
(**A)** Pavement and **(B)** guard cell numbers of lines indicated in the graph. Averages ± SEM of three biological repeats using 3 plants per line for each repeat are shown (n = 9). For statistical analysis, one-way ANOVA with Tukey’s multiple comparison was done, significant differences to the wildtype Col-0 are shown (*** p ≤ 0.001).

**Supplemental Figure 3.**
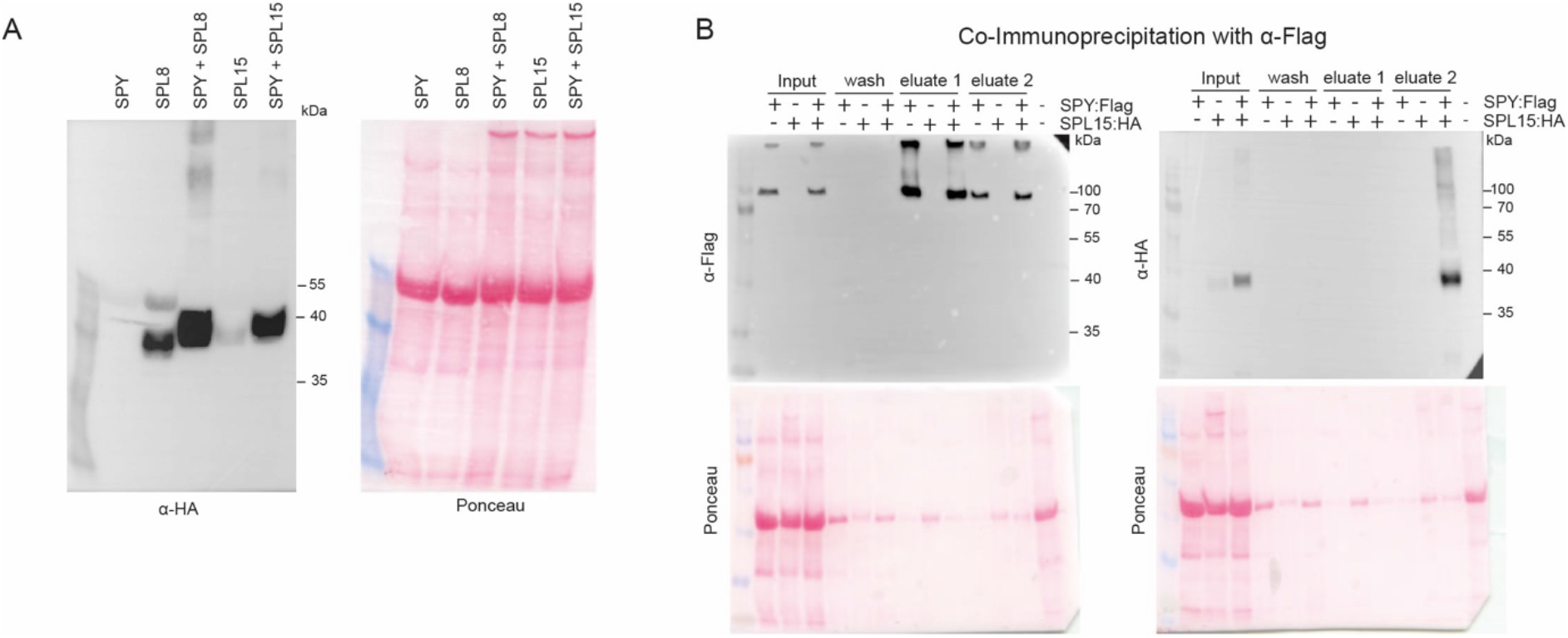
Full membranes of blots shown in Figure 7.

## Notes

### Competing Interest Statement

The authors have declared no competing interest.

### Summary of Updates

For this revision, we added phenotyping data contributed by Alexandra Baekelandt and Dirk Inze. In additional experiments, they analysed the interaction of O-glycosylation and SPLs during leaf and rosette growth by measuring leaf- and rosette size, leaf growth rates and cellular analysis. Furthermore, we included additional data on flowering of 35S::miRNA156a in O-glycosylation mutant backgrounds.

